# Extensive proteomic and transcriptomic changes quench the TCR/CD3 activation signal of latently HIV-1 infected T cells

**DOI:** 10.1101/2020.06.29.177394

**Authors:** Eric Carlin, Braxton Greer, Alexandra Duverger, Frederic Wagner, David Moylan, Alexander Dalecki, Shekwonya Samuel, Mildred Perez, Kelsey Lowman, Steffanie Sabbaj, Olaf Kutsch

## Abstract

Although the ability of HIV-1 to reside in a latent state in CD4+ T cells constitutes a critical hurdle to a curative therapy, the biomolecular mechanisms by which latent HIV-1 infection is established and maintained are only partially understood. E*x vivo* studies have shown that T cell receptor/CD3 stimulation only triggered HIV-1 reactivation in a fraction of the latently infected CD4+ T cell reservoir, suggesting that parts of the T cell population hosting latent HIV-1 infection events are altered to be TCR/CD3-activation-inert. We provide experimental evidence that HIV-1 infection of primary T cells and T cell lines indeed generates a substantial amount of TCR/CD3 activation-inert latently infected T cells. HIV-1 induced host cell TCR/CD3 inertness is thus a conserved mechanism that contributes to the stability of latent HIV-1 infection. Proteomic and genome-wide RNA-level analysis comparing CD3-responsive and CD3-inert latently HIV-1 infected T cells, followed by software-based integration of the data into protein-protein interaction networks (PINs) suggested two phenomena to govern CD3-inertness: (i) the presence of extensive transcriptomic noise that affected the efficacy of CD3 signaling and (ii) defined changes to specific signaling pathways. Validation experiments demonstrated that compounds known to increase transcriptomic noise further diminished the ability of TCR/CD3 stimulation to trigger HIV-1 reactivation. Conversely, targeting specific central nodes in the generated PINs such as STAT3 improved the ability of TCR/CD3 activation to trigger HIV-1 reactivation in T cell lines and primary T cells. The data emphasize that latent HIV-1 infection is largely the result of extensive, stable biomolecular changes to the signaling network of the host T cells harboring latent HIV-1 infection events. In extension, the data imply that therapeutic restoration of host cell TCR/CD3 responsiveness could enable gradual reservoir depletion without the need for therapeutic activators, driven by cognate antigen recognition.

**AUTHOR SUMMARY:** A curative therapy for HIV-1 infection will at least require the eradication of a small pool of CD4+ helper T cells in which the virus can persist in a latent state, even after years of successful antiretroviral therapy. It has been assumed that activation of these viral reservoir T cells will also reactivate the latent virus, which is a prerequisite for the destruction of these cells. Remarkably, this is not the case and following application of even the most potent stimuli that activate normal T cells through their T cell receptor, a large portion of the latent virus pool remains in a dormant state. Herein we demonstrate that a large part of latent HIV-1 infection events reside in T cells that have been rendered activation inert by the actual infection event. We provide a systemwide, biomolecular description of the changes that render latently HIV-1 infected T cells activation inert and using this description, devise pharmacologic interference strategies that render initially activation inert T cells responsive to stimulation. This in turn allows for efficient triggering of HIV-1 reactivation in a large part of the latently HIV-1 infected T cell reservoir.

## INTRODUCTION

Antiretroviral therapy (ART) efficiently suppresses HIV-1 replication below detection levels of diagnostic assays, but ART does not eliminate latent viral reservoirs, enabling HIV-1 to persist for the lifetime of a patient and rebound whenever ART is interrupted [1]. The most comprehensive evidence for viral persistence has been presented for latent HIV-1 infection events residing in long-lived, resting CD4 memory T cells [2-4]. Intuitively, this would explain the stability of the latent HIV-1 reservoir, as T cell memory can persist for the life-time of an individual. However, while immunological memory can persist for a life-time, individual memory T cells are relatively short lived. The half-life of individual CD4^+^ central memory T cells (T_CM_ cells) that are thought to serve as host cells of latent HIV-1 infection events ranges between 20 and ∼100 days and is generally shorter in HIV patients than in healthy individuals, with most studies suggesting a half-life τ_1/2_ <50 days [5-8]. With an assumed half-life of τ_1/2_ = 50 days and an initial reservoir consisting of 1×10^6^ latently HIV-1 infected CD4^+^ T_CM_ cells, it should take less than three years after the onset of ART for the last latently infected T_CM_ cell to disappear. This is obviously not the case and evidence has been provided that preferential or homeostatic proliferation of latently HIV-1 infected T cells in the absence of reactivation can contribute to the stability of the latent reservoir [9, 10]. While homeostatic T cell proliferation must contribute to the stability of the latent HIV-1 reservoir, it is only one contributor to lifelong immunological memory. The second component, repeated exposure to cognate antigen and subsequent re-expansion of the specific memory T cells are also crucial components of lifelong memory maintenance. How the latent HIV-1 infection pool in the memory CD4^+^ T cell population remains stable despite the expected and likely required exposure of latently HIV-1 infected T cells to their cognate antigen, and the resulting potent biological activation of the host T cells, has not been detailed. Over time, encounters with cognate antigen should cause a continuous contraction of the latent HIV-1 reservoir [11]. The absence of a measurable reservoir decay despite the expected continuous encounter of cognate-antigen and subsequent T cell activation could be explained by the idea that memory T cells hosting latent HIV-1 infection events have been altered to exhibit an activation-inert phenotype. TCR/CD3 activation-inertness would explain the inability of early therapeutic interventions (e.g. IL-2 or anti-CD3 mAb OKT3) to trigger a meaningful decrease in the size of the latent HIV-1 reservoir [12-14]. An activation-inert host cell phenotype would also explain findings that a significant part of the latent HIV-1 reservoir is resistant to *ex vivo* T cell activation, in the absence of a repressive chromatin environment at the reactivation-resistant viral LTRs [15].

Due to the rarity of latently HIV-1 infected T cells *in vivo*, and specifically the inability to identify or isolate TCR/CD3 activation-inert latently HIV-1 infected T cells from patient samples, the molecular basis of this phenomenon cannot be studied in primary T cells derived from HIV patients on ART (HIV/ART patients).

Following the demonstration that TCR/CD3-inertness also occurs in population-based *in vitro* models of latent HIV-1 infection that use either primary T cells or T cell lines, we opted to describe the molecular basis of this phenomenon in these models utilizing a systems biology approach that describes CD3-inertness at the transcriptome and proteome level. RNA-seq analysis data revealed extensive changes to the gene expression signature of latently infected T cells, regardless of their CD3-response status. Surprisingly, network analysis software was not able to assign any specific network motif to these data, let alone recognize the specific TCR/CD3 signaling deficiency, suggesting that nonspecific transcriptomic noise maybe a hallmark of latently HIV-1 infected cells. This would be consistent with reports that used single cell RNA-seq analysis to describe high levels of heterogeneity between latently HIV-1 infected T cells [16, 17]. In contrast, network analysis based on protein-level data (kinome array analysis) that described the expression and phosphorylation status of a central set of proteins detected the impairment of the TCR/CD3 signaling pathway and identified specific targets that could be pharmacologically manipulated to largely restore TCR/CD3 induced HIV-1 reactivation. Based on these data we discuss the role of both, transcriptomic noise and stable protein level changes of the host cell signaling network for the control of latent HIV-1 infection.

## RESULTS

### Impairment of the TCR/CD3 activation response in T cells from HIV/ART patients

As a result of the scarcity of latently HIV-1 infected T cells *in vivo*, it is evident that a comprehensive description of the molecular basis of TCR/CD3 inertness, in particular at the protein level cannot be done using primary T cell material from HIV/ART patients. Nevertheless, to ensure relevance of any such effort in other model systems of latent HIV-1 infection, it would be important to guide such efforts by data that are at least derived from a primary T cell population that acts as a relevant experimental surrogate system. Thus, we initially sought to identify T cell populations in samples from HIV/ART patients that would exhibit an impaired response to TCR/CD3 stimulation. Given the substantial body of literature describing an impaired ability of T cells from HIV/ART patients to respond to stimulation, we reasoned that anti-CD3/CD28 mAb-induced proliferation would allow for the identification of a TCR/CD3 activation impaired surrogate T cell subpopulation [18-26].

As we compared the proliferation response of T cells from healthy control donors and HIV/ART patients, we were surprised that impairment of cell proliferation was not limited to a small subpopulation of T cells, but rather was observed for the majority of CD4+ T cells from HIV/ART patients. Following CD3/CD28 mAb stimulation CD4+ T cells from most healthy control donors (8 of 11) showed a coordinated, discrete proliferation response profile. In contrast, the proliferation response of T cells from the majority of tested virally suppressed HIV/ART patients (8 of 12) was not necessarily quantitatively reduced, but qualitatively impaired, uncoordinated and asynchronous (Figure 1A; Supplemental Figure 1). These data suggested that a large portion of the T cells from patients would have an altered biomolecular phenotype that seemed to affect the TCR/CD3 stimulation induced proliferation response. Given the global nature of this phenotype, we decided to forgo any enrichment of specific T cell subpopulations and use unsorted T cell populations for the initial systems biology analysis.

**Figure 1:**
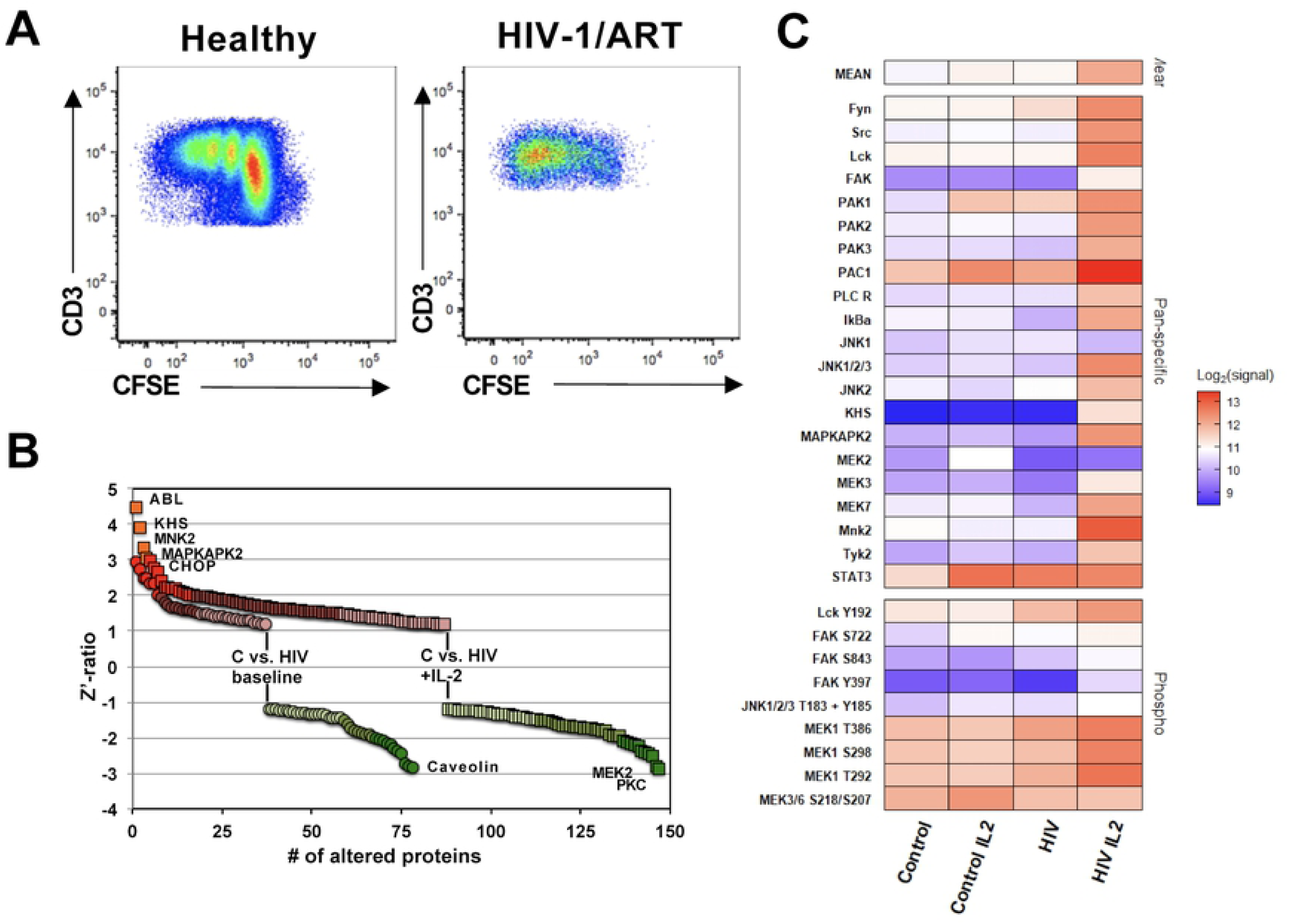
T cells from HIV-1 patients on ART exhibit altered activation response phenotypes. (**A**) T cells from healthy individuals and HIV/ART patients were stained with CFSE and stimulated with a CD3/CD28 mAb combination. The proliferative response of the T cells, measured as decrease in CFSE fluorescence intensity, was determined on day 4 post stimulation using flow cytometric analysis. The coordinated CFSE profile shown is representative for a total 8 of 11 healthy controls, whereas an asynchronous CFSE profile was found for 8 of 11 HIV/ART patients. (**B**) Kinome array analysis was performed using T cells from 9 healthy individuals and 8 HIV/ART patients. Samples were produced either at baseline or following 24h of IL-2 incubation. The samples for each experimental conditions were pooled to minimize individual variations and loaded onto Kinexus antibody KAM-850 arrays and expression levels and phosphorylation states of a total of 510 kinases and phosphatases were determined. Signals that were altered in samples from HIV/ART patients relative to healthy controls at baseline or following IL-2 stimulation were plotted. (**C**) Focused analysis of kinase and transcription factor signals from the kinome array involved in the TCR/CD3 signaling pathway represented as a heat map (expression or phosphorylation) for each of the four experimental conditions.

As proteins are the ultimate mediators of the TCR/CD3 signaling pathway, we hypothesized that data describing the altered proteomic state of T cells of patients could help to prioritize data describing the altered state of latently HIV-1 infected T cells. Also, the TCR/CD3 signal is generally not regulated or transmitted through gene regulation, but by protein phosphorylation effects. Accordingly, we used Kinexus antibody arrays that interrogate significant parts of the proteome at the level of protein expression and protein phosphorylation. We chose two experimental conditions, (i) unstimulated and (ii) overnight exposure to IL-2, a simple experimental surrogate of conditions that T cells would encounter in an immunologically more active environment such as lymph nodes. More than 600 signals were found higher than the predetermined intensity cut-off. Z-ratios were used to identify signals that differed between two conditions. At baseline, the kinome array experiments identified a total of 78 differences (37 up-/41 down-regulated) between T cells from healthy controls and HIV/ART patients, but 147 differences (87 up/60 down) following overnight exposure to IL-2, suggesting that T cells from HIV/ART patients are hyper-responsive (Figure 1B). A focused analysis of the data for significantly altered proteins that have been reported to be involved in TCR/CD3 signaling demonstrated the impairment of this pathway in T cells from HIV/ART patients, in particular following IL-2 exposure, but also at baseline (Figure 1C).

Ranking the importance of proteins within a signaling network by the magnitude of the observed signal changes would bias the analysis against the possible contribution of low amplitude signals. Instead we used a direct interaction algorithm from the MetaCore network analysis software to generate protein-protein interaction networks that exclusively use the identified, altered proteins (seed nodes). The resulting network allows to prioritize proteins based on their functional importance as indicated by the numbers of interactions within the network, rather than just signal amplitude. A total of 83% of the 230 seed nodes were integrated in the direct interaction network (Supplemental Figure 2), a sign of the quality of the generated data, as we have previously demonstrated that a random list of genes is not linked by this algorithm [27]. 54 nodes had more than 20 interactions in the network, suggesting an overall high degree of linkage between the nodes and a relatively flat signaling hierarchy. Within this network the central proteins (hubs) were p53 (108 interactions), STAT3 (93 interactions), Src (73 interactions), c-Jun (68 interactions), and ß-catenin (67 interactions), all factors that play key roles in TCR/CD3 signaling (Table 1) [28-35]. Consistent with the observed functional phenotype of T cells from HIV/ART patients that indicates an impairment of cell cycle activity following TCR/CD3 stimulation (Figure 1A), as well as the experimental conditions (IL-2 stimulation), network enrichment analysis ranked G1-S growth factor regulation (p=9.91E-25), IL-2 signaling (p= 1.26E-21) and TCR signaling (p= 3.60E-19) in the top five altered network motifs (Table 2). Primarily, these data validate the ability of kinome analysis in conjunction with MetaCore networking software to accurately detect and describe experimental conditions (IL-2 signaling) and phenotypes (TCR signaling, cell cycle impairment). The data suggest that permanent changes to the signaling network of T cells obtained from HIV/ART patients, may not be limited to latently HIV-1 infected T cells. Under the assumption that latently infected T cells may only represent an extreme phenotype of a T cell dysfunctionality spectrum, possibly a long-term consequence of the initial infection and the widespread destruction of the immune system during the initial infection, the kinomic data describing the altered molecular phenotype of primary T cells from HIV/ART patients should provide an efficient tool to distinguish between HIV-1 infection induced changes that are shared by all T cells (primary and immortalized), and changes that are limited to latently HIV-1 infected T cell lines.

**Table 1:** Top ranked network nodes from protein-protein interaction network describing the different signaling modalities of T cells from healthy individuals and HIV patients.

| Rank | Name | Number of Interactions |
| --- | --- | --- |
| 1 | p53 | 108 |
| 2 | STAT3 | 93 |
| 3 | Src | 73 |
| 4 | c-Jun | 68 |
| 5 | Beta-catenin | 67 |
| 6 | Caspase-3 | 65 |
| 7 | CDK1 (p34) | 59 |
| 8 | AKT1 | 58 |
| 9 | STAT1 | 52 |
| 10 | p27KIP1 | 51 |
| 11 | EGFR | 48 |
| 12 | PKC-delta | 45 |
| 13 | c-Raf-1 | 43 |
| 14 | c-Abl | 41 |
| 15 | c-Fos | 40 |
| 16 | ERK1 (MAPK3) | 40 |
| 17 | CDK2 | 37 |
| 18 | CDK5 | 37 |
| 19 | PKC-alpha | 37 |
| 20 | Rb protein | 35 |

**Table 2:** Network enrichment analysis using kinome data describing the different signaling modalities of T cells from healthy donors and HIV patients.

| Rank | Networks | p-value |
| --- | --- | --- |
| 1 | G1-S Growth factor regulation | 9.951E-25 |
| 2 | Hemopoiesis, Erythropoietin pathway | 1.202E-22 |
| 3 | Anti-Apoptosis (MAPK and JAK/STAT) | 7.410E-22 |
| 4 | IL-2 signaling | 1.267E-21 |
| 5 | TCR signaling | 3.608E-19 |
| 6 | G1-S Interleukin regulation | 5.718E-19 |
| 7 | NOTCH signaling | 9.348E-19 |
| 8 | Regulation of initiation | 5.627E-18 |
| 9 | Lymphocyte proliferation | 1.034E-16 |
| 10 | Apoptosis stimulation by external signals | 1.686E-16 |

### HIV-1 infection generates a TCR/CD3 activation inert population of latently infected T cells

Next, we sought to confirm previous reports that a fraction of the latently HIV-1 infected primary T cell population would be inert to TCR/CD3 stimulation, at least regarding the ability of this stimulus to trigger HIV-1 reactivation [15, 36]. For this purpose we used a modification of a well-established primary T cell model of HIV-1 latency [36-42]. Briefly, activated CD4+ T cells were infected with a HIV-1 reporter virus and RT inhibitors were added beginning 72hr post infection. At four weeks post infection, the infection cultures were sorted to remove all actively infected and therefore GFP-positive T cells to lower the signal background. The CD4+T cells were then left unstimulated, stimulated with anti-CD3/CD28 mAb coated beads (ImmunoCult) or stimulated with PMA/ionomycin. The latter is an experimental means to trigger potent activation and proliferation in primary T cells while completely bypassing the TCR/CD3/CD28 pathway. 72hr post activation the samples were subjected to flow cytometric analysis. As stimulation was performed in the presence of RT inhibitors, an increase in the frequency of GFP-positive T cells over background infection in the control sample would reflect reactivated, previously latent HIV-1 infection events. In line with the idea that a fraction of the host T cells of latent HIV-1 infection events have been rendered inert to TCR/CD3 activation, stimulation of CD4+ T cell infection cultures generated by using T cell material from three different donors with a highly potent PMA/ionomycin combination consistently triggered higher levels of HIV-1 reactivation levels than antibody mediated CD3/CD28 stimulation (Figure 2A).

**Figure 2:**
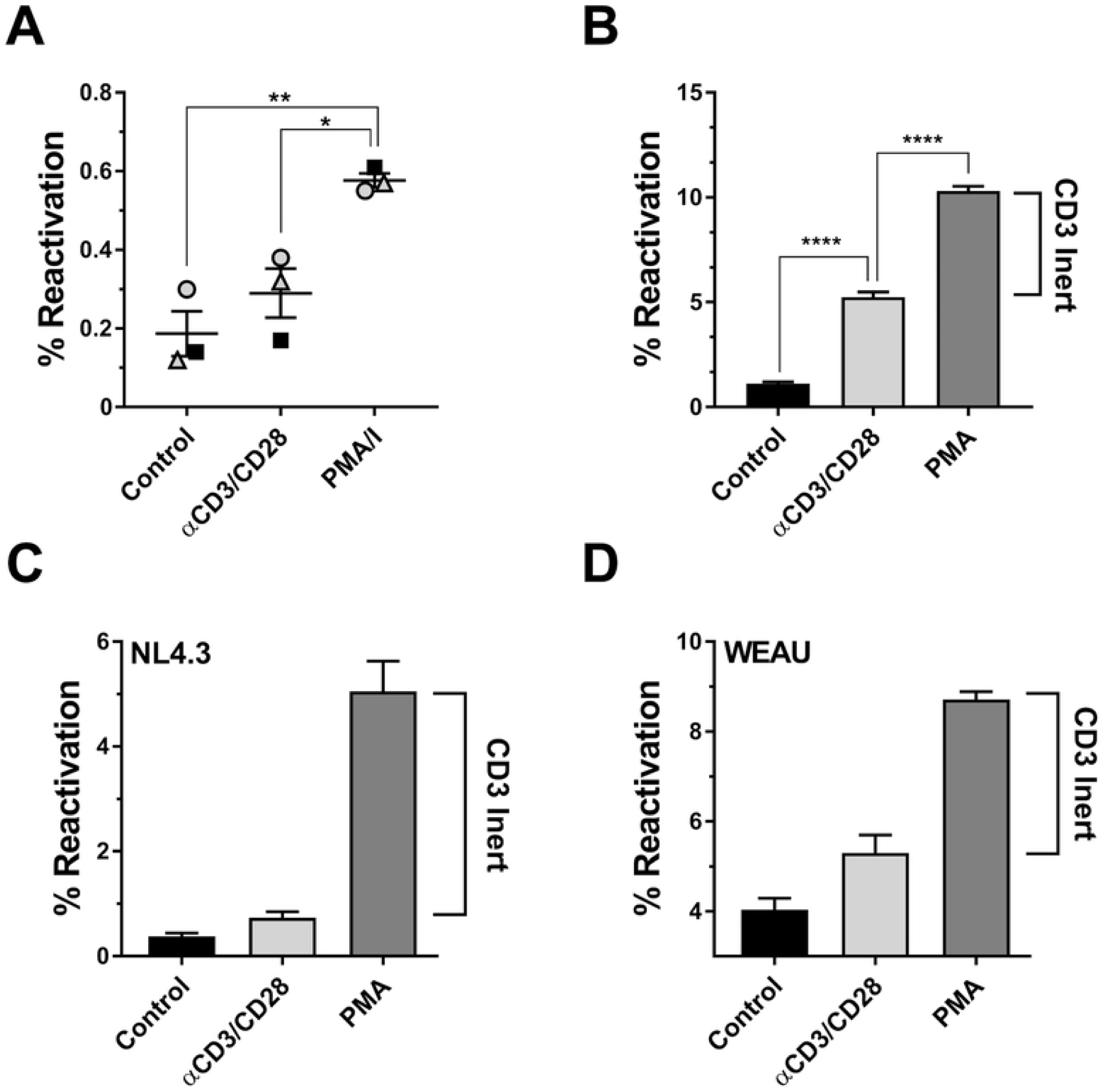
HIV-1 infection induced TCR/CD3 activation inertness is conserved between primary T cells and T cell lines. (**A**) Primary CD4+ T-cells from three donors were activated, infected with a HIV-GFP reporter virus and 4 weeks post infection, residual GFP-positive, and therefore actively infected T cells were removed by cell sorting. The T cell cultures were then left unstimulated (control), stimulated with anti-CD3/CD28 mAb coated beads or stimulated with PMA/ionomycin. Using GFP expression as a surrogate marker, levels of active/reactivated HIV-1 expression were determined 72 hours post stimulation using flow cytometric analysis for GFP expression. (**B**) Jurkat T cells were infected with a HIV-1 NL4-3 based GFP reporter virus as previously described [49, 50]. The T cell populations were then cultured for a total of 60 days in the presence of RT inhibitors to prevent de novo infection. At this time, samples of the infection cultures were either left unstimulated, treated with anti-CD3/CD28 mAb coated beads or stimulated with PMA. Baseline HIV-1 expression levels in the untreated cells and HIV-1 reactivation in the stimulated cultures were determined using flow cytometric analysis for GFP expression. Similar experiments were performed using a reporter T cell line that expresses GFP when actively HIV-1 infected (J-R5D4 cells). J-R5D4 T cells were infected with (**C**) HIV-1 NL4-3 or (**D**) HIV-1 WEAU, a primary HIV-1 patient isolate. Following six weeks of culture in the presence of RT inhibitors reactivation experiments were performed as in (B) and reactivation levels were determined by flow cytometric analysis for GFP expression. Data plotted as mean ± standard deviation and the significance of the response differences were determined by one-way ANOVA with multiple comparisons.

While these results confirm that CD3-inertness could be a contributing factor for the stability of the latent HIV-1 reservoir, the primary T cell model of HIV-1 latency is not conducive to comprehensive high resolution systems biology analysis of biomolecular changes in the signaling networks of latently HIV-1 infected T cells. The relatively small number of latently HIV-1 infected T cells that can be generated in the model limits analysis to single cell RNA-seq. Protein level analysis which is likely central to the identification of pertinent blocks in the signaling pathways at the phosphorylation level would be completely impractical due to the large amount of required cell material. As for primary T cell models from patients, the relatively small number of latently HIV-1 infected T cells in these populations can only be identified after a stimulus that triggers HIV-1 reactivation is provided, which (i) still would not allow to discover activation inert subpopulations and (ii) make studies of the baseline signaling networks impossible.

Given that Jurkat T cells, despite some shortcomings, were used to decipher the fundamental molecular biology of TCR/CD3 signaling, the effect of CD3 activation on HIV-1 replication [43-46] and are the most commonly used CD4+ T cell line to study HIV-1 latency [47-49], we next explored whether the phenomenon of HIV-1 infection induced CD3-inertness would also occur in HIV-1 infection cultures of Jurkat T cells. Similar to previous efforts made to generate latently HIV-1 infected reporter T cell clones, we infected Jurkat T cells with a HIV-1 reporter virus and followed the establishment of latent HIV-1 infection at the population level [27, 38, 39, 48-50]. After 6 weeks of culture in the presence of RT inhibitors, active background infection was reduced to <2% (Control) and addition of PMA triggered reactivation in ∼10% of the T cell population (Figure 2B). Antibody mediated stimulation of CD3/CD28 only triggered HIV-1 reactivation in ∼5% of the cells. This phenomenon could be reproduced in reporter T cell populations (J-R5D4) that were infected with either the recombinant clone HIV-1 NL43 (Figure 2C) or with HIV-1 WEAU, a primary R5-tropic patient isolate (Figure 2D) [50, 51], where PMA induced 10-fold or 3-fold more reactivation, respectively, than CD3/CD28 stimulation. As described for primary T cells, in all experiments only a fraction of latent HIV-1 infection events that could be reactivated by PMA stimulation would respond to CD3/CD28 stimulation. These data suggest that HIV-1 infection induced CD3-inertness is a fundamental phenomenon that is shared between primary T cells and T cell lines.

### Generation of CD3-responsive and CD3-inert latently HIV-1 infected T cell clones

While the percentage of latently HIV-1 infected T cells in long-term HIV-1 infection cultures based on Jurkat T cells (Figures 2B-D) is higher than in *in vitro* models of latent HIV-1 infection in primary T cells (Figure 2A), these T cell populations are a mixture of cells that are not infected, infected with defective viruses and 5-10% latently HIV-1 infected T cells, likely representing a spectrum of CD3 response phenotypes. The signaling noise resulting from the cellular diversity would render these T cell populations suboptimal to perform detailed systems biology-based analysis of the mechanisms governing CD3-inertness. We thus generated two latently HIV-1 infected T cell clones that would represent CD3-responsive (JWEAU-A10) and CD3-inert (JWEAU-C6) T cell phenotypes (Figure 3A), despite comparable level of CD3 or CD28 expression (Figure 3B). PMA triggered similar HIV-1 reactivation response levels in both T cell clones (Figure 3A), but even saturating amounts of CD3/CD28 mAbs only triggered reactivation in JWEAU-A10 T cells, but not in JWEAU-C6 cells. (Figure 3B). CD3-inertness of JWEAU-C6 T cells was not a function of the utilized antibody clone (OKT3, UCHT1, HIT3A) (Figure 3E). Using a TransAM assay to quantify relative NF-κB activity over time we could show that TCR/CD3 stimulation of JWEAU-A10 T cells produced a strong NF-κB response that became apparent as early as 15 minutes after stimulation, peaked after one hour and was maintained at an elevated level for >4 hours. Throughout the same observation period, TCR/CD3 stimulation would not trigger an efficient NF-κB signal in JWEAU-C6 cells (Figure 3F). This is likely the result of a block in the upstream signaling cascade and not the result of a dysfunctional canonical NF-κB pathway, as not only PMA, but also TNF-α, which triggers the NF-κB activation through a different upstream pathway would induce efficient HIV-1 reactivation in JWEAU-A10 and JWEAU-C6 T cells (Supplemental Figure 3). Of note, TLR5 signaling which triggers NF-κB activation following stimulation with bacterial flagellin was also partially compromised in JWEAU-C6 T cells when compared to JWEAU-A10 T cells (Figure 3G). This was also observed at the population level, where flagellin stimulation only triggered HIV-1 reactivation in a fraction of the latently HIV-1 infected T cell population (Figure 3H). Changes to the host cell signaling network that were induced during HIV-1 infection thus may not be limited to the TCR/CD3 signaling pathway and extend to other activating receptors/pathways that are currently investigated as targets of latency reversing agents (LRA) [52-56].

**Figure 3:**
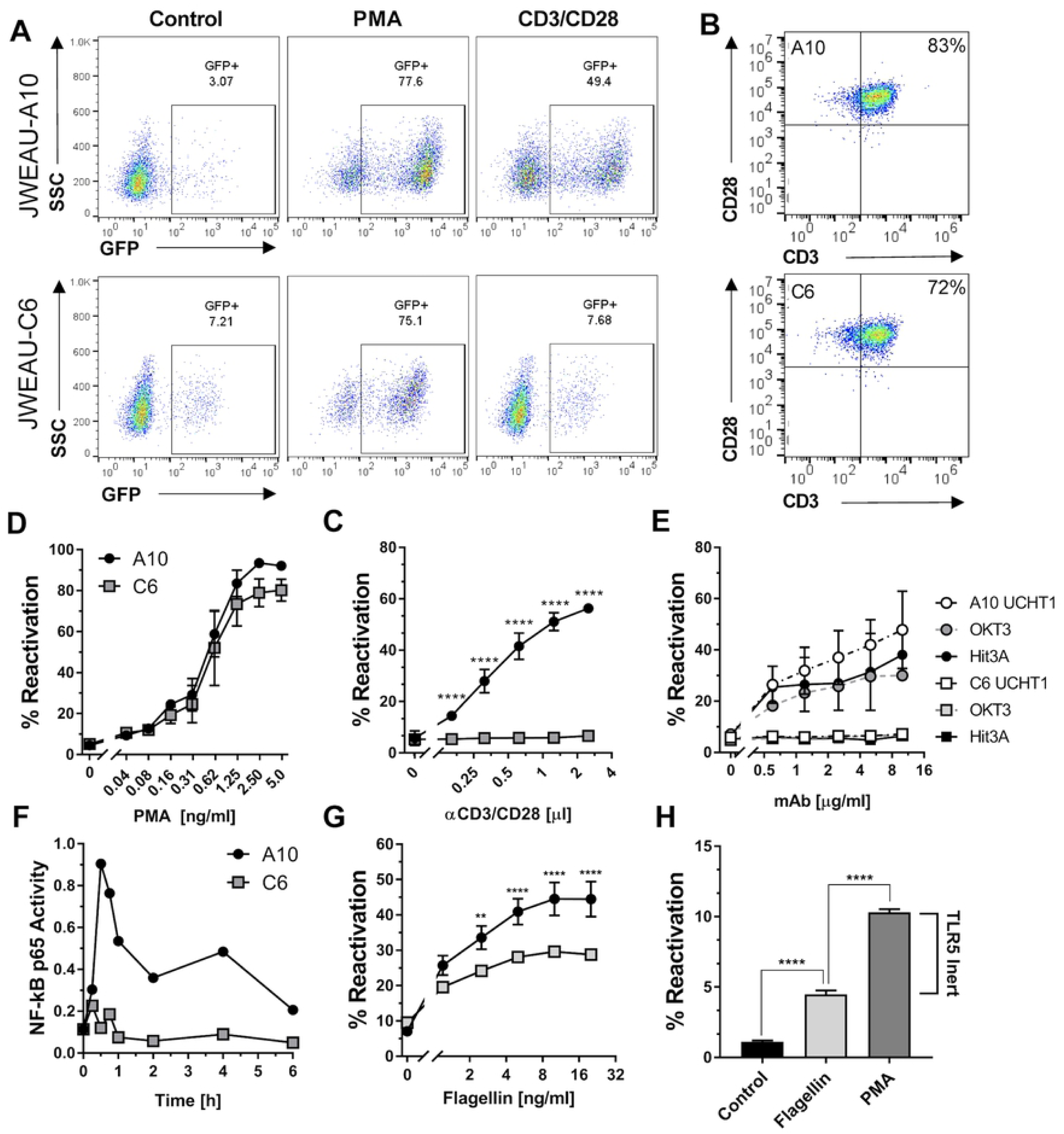
Generation of TCR/CD3-inert and -responsive latently HIV-1 infected T cell clones. (**A**) Representative flow cytometric dot plot analysis of GFP expression as a surrogate marker of HIV-1 expression in the TCR/CD3-responsive JWEAU-A10 T cells and the TCR/CD3-inert JWEAU-C6 T cells at baseline (control) and following activation by α-CD3/CD28 mAb or PMA. (**B**) Expression of CD3 and CD28 on JWEAU-A10 and JWEAU-C6 T cells as determined by flow cytometric analysis. (**C**) HIV-1 reactivation levels in JWEAU-A10 and JWEAU-C6 T cells following stimulation with increasing concentrations of the PKC activating phorbolester PMA as determined by flow cytometric analysis for GFP expression. (**D**) HIV-1 reactivation levels in JWEAU-A10 and JWEAU-C6 T cells following stimulation with increasing concentrations of α-CD3/CD28 mAb as determined by flow cytometric analysis for GFP expression. (**E**) HIV-1 reactivation levels in JWEAU-A10 and JWEAU-C6 T cells following stimulation with increasing concentrations of different α-CD3 mAbs (OKT3, UCHT1, HIT3A) as determined by flow cytometric analysis for GFP expression. (**F**) Kinetic NF-κB p65 response in JWEAU-A10 and JWEAU-C6 T cells following stimulation with α-CD3/CD28 mAb. (**G**) HIV-1 reactivation levels in JWEAU-A10 and JWEAU-C6 T cells following stimulation with increasing concentrations of the TLR5 agonist flagellin. (**H**) HIV-1 reactivation levels in JWEAU-A10 and JWEAU-C6 T cells following stimulation with increasing concentrations of flagellin. Where indicated, data represent the mean ± standard deviation of at least three independent experiments. The significance of differences between experimental conditions was determined by one-way ANOVA with multiple comparisons.

### Gene expression patterns associated with CD3-inert latently HIV-1 infected T cells

At the time, studies that have described the phenotype of latently HV-1 infected T cells have focused on changes to the transcriptome [16, 17]. To possibly align our data with these data sets, we first generated RNA-seq data describing the gene expression signatures of uninfected T cells, the CD3 activation-inert JWEAU-C6 T cells and the CD3-responsive JWEAU-A10 T cells.

A total of 7,151 genes (of 20,830 genes with any reads) were differentially expressed (likelihood ratio test, adjusted *P*-value < 0.01) across the three cell types. More stringent pairwise comparison between the parental Jurkat T cells and each of the latently HIV-1 infected T cell clones was performed with a padj < 0.01, a fold-change ≥2; and the signal for at least one the cell types had to be >250 reads. Relative to Jurkat T cells 481 genes were upregulated and 627 were downregulated in JWEAU-C6 T cells (total of 1108 genes); 537 genes were upregulated and 484 genes were downregulated in JWEAU-A10 T cells (total of 1021 genes) (Figure 4A). 354 of the downregulated and 304 of the upregulated genes were shared between the two latently HIV-1 infected T cells clones. This would mean that of the altered genes, only 45% are shared by the two latently HIV-1 infected T cell clones. In a direct comparison of the two latently HIV-1 infected T cell clones, relative to JWEAU-A10 T cells, 348 genes were downregulated and 125 were upregulated in JWEAU-C6 T cells (Figure 4B). These findings are consistent with recent reports suggesting vast heterogeneity between individual latently HIV-1 infected T cells and emphasize that HIV-1 latency can be maintained in extensively different cellular environments [16, 17]. A direct interaction network generated using the list of genes found to be differentially regulated between JWEAU-C6 and JWEAU-A10 T cells only integrated 53% of the seed nodes, and of these, 50% were connected with only a single link (Supplemental Figure 4). In average each node of the network had only three interactions. Both the failure to efficiently link all signals into the network and the low level of interactions per node suggest that a large portion of the observed gene expression changes are not functionally related and represent transcriptomic noise. In line with transcriptomic noise being the major component of the observed signal changes, motifs suggested by network enrichment analysis had very low statistical significance and, most importantly, completely failed to recognize the defined experimental phenotype, TCR/CD3 inertness (Supplemental Table 1).

**Figure 4:**
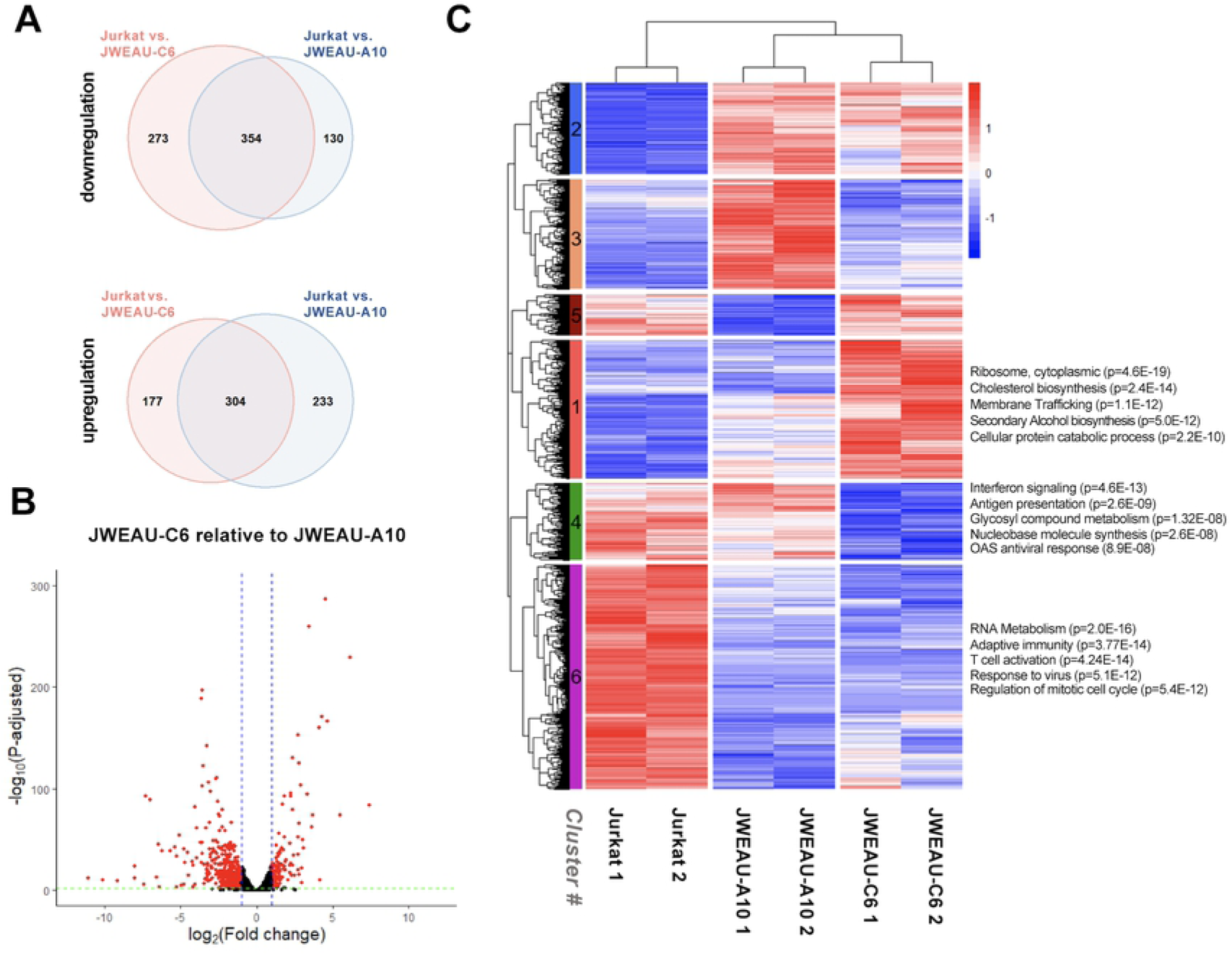
Transcriptomic analysis reveals extensive differences between uninfected and latently HIV-1 infected T cells. (**A**) Pairwise comparison of altered gene expression patterns between the parental Jurkat T cells and each of the latently HIV-1 infected T cell clones was performed with a padj < 0.01, a fold-change ≥2; and the signal for at least one the cell types had to be >250 reads. (**B**) Pairwise comparison of differentially expressed genes between JWEAU-A10 T cells and JWEAU-C6 T cells. Gene expression changes are plotted as log2 fold change in JWEAU-C6 T cells relative to gene expression levels in JWEAU-A10 T cells. (**C**) RNA-seq analysis was used to describe the genome-wide RNA expression signature of uninfected Jurkat T cells, the TCR/CD3 responsive JWEAU-A10 and the TCR/CD3-inert JWEAU-C6 T cells. For each T cell line, RNA-seq data were obtained for low and high cell density conditions. 7,151 genes with differential expression (likelihood ratio test, adjusted *P*-value < 0.01) across the three cell types were subjected to hierarchical clustering. Subtrees representing genes that were lowly or highly expressed throughout all three cells were automatically omitted through the selection criteria.

If transcriptomic noise were to mask functional motifs in the data set, it stands to reason that hierarchical clustering of the gene expression data could improve the likelihood of discovering motifs that control latency, assuming the phenomenon is driven primarily by changes to gene expression (Figure 4C). This was indeed the case. Regulation of genes in two of the sub-clusters was largely shared between the two latently HIV-1 infected T cell lines, but different from the parental T cells. Genes in cluster #2 (960 genes) and cluster #6 (2353 genes) were upregulated or downregulated, respectively, in both latently infected T cell clones when compared to uninfected T cells. This would suggest that these changes were likely common features of latency control. Gene Ontology (GO) enrichment (Figure 4C; Supplemental Figure 5) of cluster #6 genes (downregulated in both latently HIV-1 infected T cell clones) suggested changes to RNA metabolism, adaptive immune system function, the antiviral response, regulation of the mitotic cell cycle, and T cell activation. Upregulated genes (cluster #2) were found to be reported as critical for lipid metabolism, regulation of cellular component size, and actin filament-based processes. These findings are in agreement with previous descriptions of gene regulation in latently HIV-1 infected cells, suggesting that our model system recapitulates established transcriptional hallmarks of latently infected host cell changes [42, 57-91].

In the remaining four clusters, gene regulation differed between the two latently HIV-1 infected T cells, but one or the other had gene regulation patterns similar to the parental T cells (cluster #1 (1449 genes), #3 (1155 genes), #4 (805 genes), #5 (429 genes)), emphasizing that latently HIV-1 infected T cells not only differ in their gene expression signature from uninfected T cells, but also greatly differ from each other. As uninfected Jurkat T cells are known to be responsive to TCR/CD3 stimulation, we would speculate that cluster #1 and #4, in which Jurkat T cells and the responsive JWEAU-A10 T cells share expression patterns, but JWEAU-C6 gene expression patterns differ, should hold the dysregulated genes that are responsible for the TCR/CD3-inertness of JWEAU-C6 T cells. Surprisingly, none of the enriched GO terms for these genes suggested critical relevance to CD3-responsiveness. Enrichment for Gene Ontology (GO) terms using the genes in clusters #1 and #4 failed to identify GO terms associated with TCR signaling (Figure 4C and Supplemental Figure 5). We thus refocused on the genes in cluster #6 that would be descriptive of the T cell activation motif (GO term 0042110), however, detailed analysis of these data emphasized that the TCR/CD3 signaling pathway is generally impaired in latently HIV-1 infected T cells, but did not reveal relevant differences between the two latently HIV-1 infected T cell clones that could immediately explain the completely CD3-inert phenotype of JWEAU-C6 T cells (Supplemental Figure 6).

The data confirm that host T cells of latent HIV-1 infection events greatly differ from uninfected T cells and demonstrate that heterogeneity between latently HIV-1 infected primary T cells is also a characteristic of latently infected T cell lines that were generated under identical experimental conditions. The data suggest that impairment of the TCR signaling pathway is a fundamental characteristic that is shared between latently infected T cell clones, but little information on potential targets that could influence the stability of latent HIV-1 infection could be directly extracted from the data set. Lastly, given the sheer magnitude of the gene expression differences that are detected in the latently HIV-1 infected T cells, and the likely resulting misalignment of many cellular functionalities, it is tempting to speculate that transcriptomic noise could induce an unspecific signaling threshold to TCR/CD3 signaling.

### Kinomic description of CD3-activation inert latently HIV-1 infected T cells

One possible explanation for the inability of the RNA-level analysis to directly identify the underlying functional impairment (CD3-inertness) that stabilizes latent HIV-1 infection is that major regulation effects occur at the level of post-transcriptional and post-translational modifications (e.g. protein phosphorylation). Changes to the proteome at the level of protein expression and phosphorylation would be the most direct indicators of functional modifications of the host cell signaling network that cause CD3-inertness and contribute to the stability of latent HIV-1 infection. To identify protein regulation effects that differ between CD3-responsive and CD3-inert T cells we again used antibody arrays. Between JWEAU-A10 and JWEAU-C6 T cells the array identified a total of 123 significant differences. Of these signals, 74% were phospho-site specific and 26% were pan-specific. This was consistent with the general distribution of pan-specific or phospho-site specific antibodies on these arrays and did not indicate any bias, however, it provides an estimate of the number of changes occurring at the level of protein phosphorylation. The MetaCore direct interaction algorithm efficiently linked these differentially regulated signals. 86% of the seed nodes were linked into a direct protein-protein interaction network, which had an average degree of 14 interactions per node, a sign that likely more than one pathway within the overall network was affected, and no single master switch controlled the recorded changes (Supplemental Figure 7). The three highest linked nodes in the network were p53 (55 interactions), STAT3 (47 interactions), and c-Src (46 interactions) (Table 3), which paralleled the findings in primary T cells from HIV/ART patients (Table 1). Also, consistent with the functional phenotype, pathway enrichment analysis identified *Immune Response/TCR signaling* (p=6.78E-14) as a highly ranked altered motif (Table 4). There was additional overlap with other motifs that had been identified as altered in primary T cell material from HIV/ART patients, suggesting that HIV-1 infection induced changes that are shared between primary T cells and T cell lines are not limited to the immediate TCR/CD3 signaling pathway (e.g. G1-S Growth Factor regulation (p=3.97E-16), NOTCH signaling(p=1.87E-16), regulation of initiation (p= 2.2E-15)).

**Table 3:**
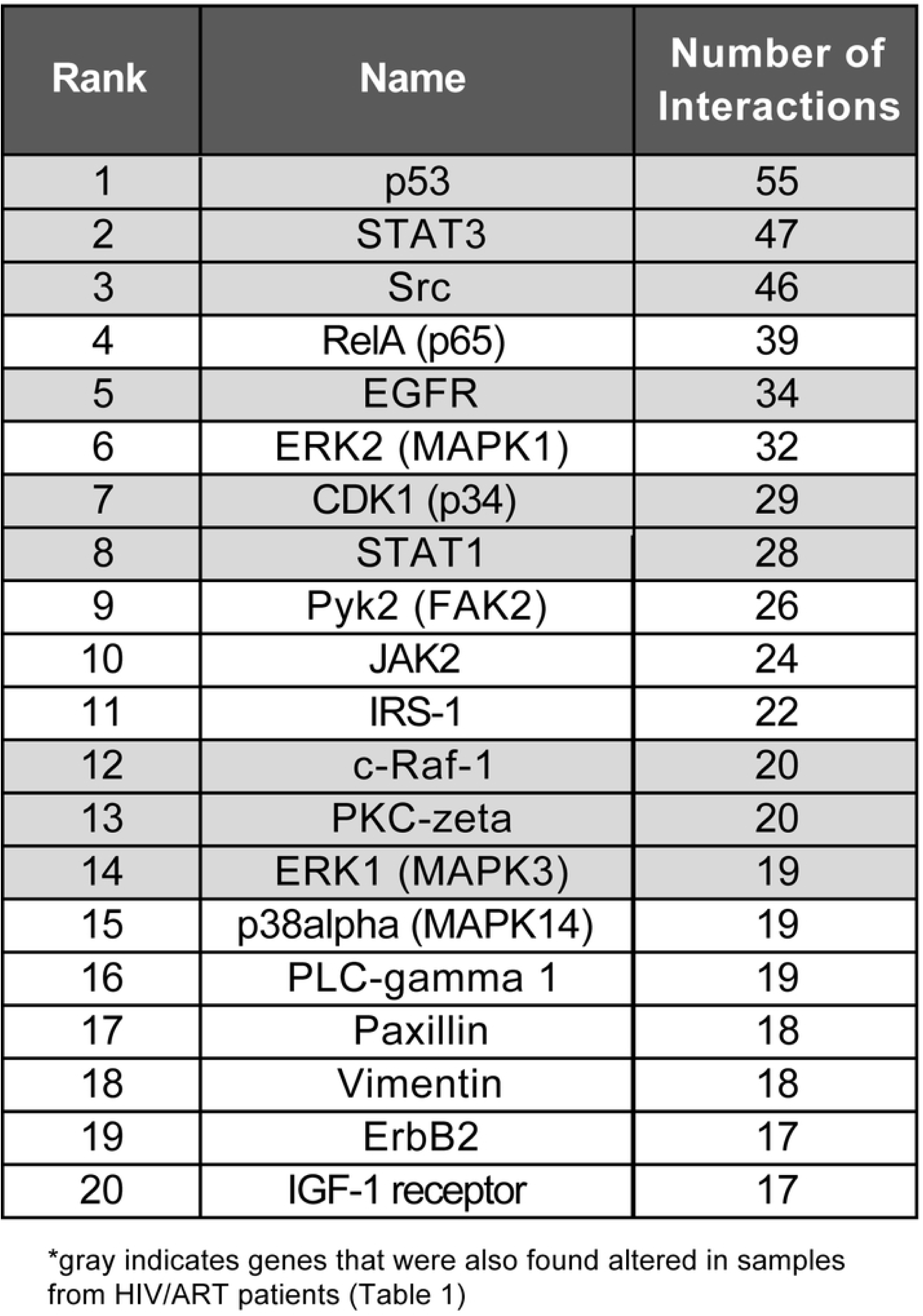
Top ranked network nodes from protein-protein interaction network describing the different signaling modalities of JWEAU-A10 and JWEAU-C6 (inert) T cells.

**Table 4:** Network enrichment analysis of kinome data derived from differentially TCR responsive cell lines identifies numerous networks observed in primary T cell proteomic data.

| Rank | Networks | p-value |
| --- | --- | --- |
| 1 | Hemopoiesis, Erythropoietin pathway | 3.101E-17 |
| 2 | NOTCH signaling | 1.874E-16 |
| 3 | G1-S Growth factor regulation | 3.716E-16 |
| 4 | Regulation of initiation | 2.196E-15 |
| 5 | ERBB-family signaling | 2.879E-15 |
| 6 | G1-S Interleukin regulation | 3.979E-14 |
| 7 | TCR signaling | 6.781E-14 |
| 8 | Positive regulation cell proliferation | 6.678E-13 |
| 9 | Regulation of epithelial-to-mesenchymal transition | 8.713E-13 |
| 10 | Anti-Apoptosis (MAPK and JAK/STAT) | 1.333E-12 |

\*gray indicates genes that were also found altered in samples from HIV/ART patients (Table 1)

### Validating the role of transcriptomic noise as reactivation threshold for TCR/CD3 signaling

The gene expression and protein level analysis suggested two potential biomolecular mechanisms that contribute to TCR/CD3 activation inertness in a large fraction of the latently HIV-1 infected T cell reservoir: (i) the presence of transcriptomic noise that could interfere with accurate signal transduction and (ii) stable changes of the activity or responsiveness of individual network nodes detected mostly at the protein level, that could act as central switches for the transmission of the TCR/CD3 signal. To test the first hypothesis, we took advantage of findings by others describing that histone deacetylase (HDAC) inhibitors, BET protein inhibitors or cell differentiating agents, through a variety of specific and off-target effects trigger genome-wide changes to gene expression patterns, and as such create transcriptomic noise [92-102]. Some have actually argued that HDAC inhibitors would trigger HIV-1 reactivation by inducing an increase in genetic noise [103] and the varying potential of different histone deacetylase inhibitors to trigger HIV-1 reactivation has been linked to the induction of differential host cell responses [104].

We reasoned that if transcriptomic noise contributes to a TCR/CD3 activation threshold in latently HIV-1 infected T cells, then treatment of the CD3-responsive JWEAU-A10 T cells with HDAC inhibitors, BET inhibitors or cell differentiating agents, would increase the level of transcriptomic noise and reduce the ability of TCR/CD3 activation to trigger HIV-1 reactivation. Following a 24 hours pretreatment period to allow each inhibitor to exert its full effect, non-toxic concentrations of the histone deacetylase inhibitor suberanilohydroxamic acid (SAHA; vorinostat) had no direct effect on HIV-1 reactivation in either, JWEAU-A10 and JWEAU-C6 T cells (Figure 5A). As predicted, SAHA then decreased the TCR/CD3 stimulation-induced HIV-1 reactivation response of JWEAU-A10 T cells, while not restoring a TCR/CD3 response in JWEAU-C6 T cells. The cell differentiating compound hexamethylene bisacetamide (HMBA), which has been reported to trigger HIV-1 reactivation in some experimental systems [105, 106], did not directly trigger HIV-1 reactivation (Figure 5B). However, in line with its ability to induce transcriptomic noise HMBA suppressed the TCR/CD3 stimulation-induced HIV-1 reactivation response of JWEAU-A10 T cells. HMBA had no effect on latent HIV-1 infection in JWEAU-C6 T cells, neither by itself nor following TCR/CD3 stimulation. Lastly, we tested the effect of JQ1, a Brd4 inhibitor that has been reported to not only affect global cellular gene expression, but to trigger low level HIV-1 reactivation [107-109]. At the utilized concentration JQ1 triggered modest levels of HIV-1 reactivation by itself in JWEAU-A10 T cells, but again suppressed the ability of TCR/CD3 stimulation to trigger HIV-1 reactivation (Figure 5C). These data provide evidence that compounds that are known to generate transcriptomic noise impair the ability of TCR/CD3 signaling to trigger HIV-1 reactivation in otherwise CD3-responsive T cells, and thereby indirectly validate a role for transcriptomic noise as a HIV-1 reactivation threshold component.

**Figure 5:**
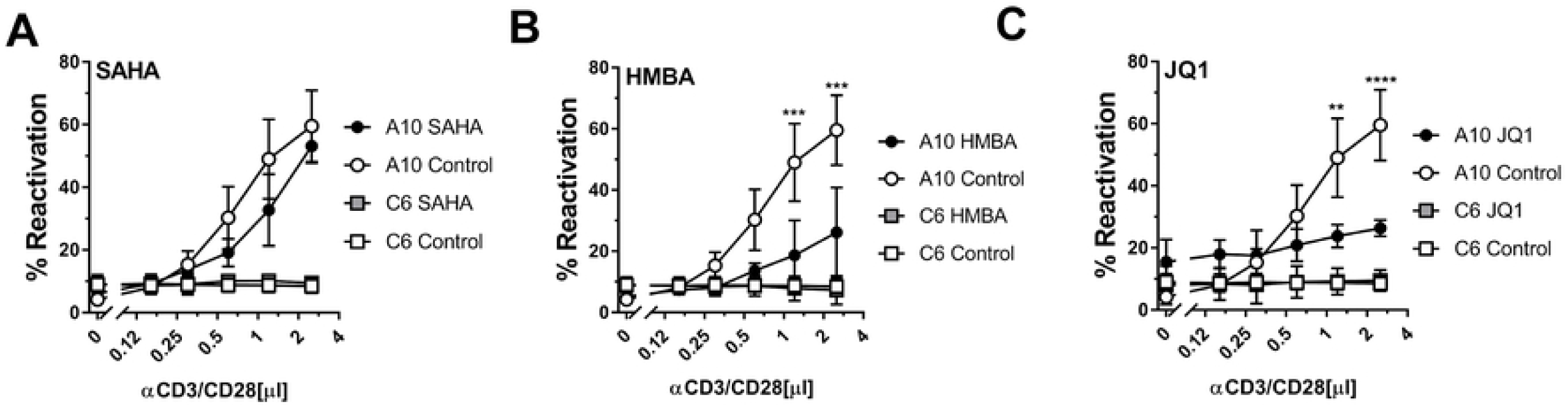
Induction of genetic noise suppresses TCR/CD3 induced HIV-1 reactivation. To determine if transcriptomic noise contributes to the stability of latent HIV-1 infection we pretreated the TCR/CD3 responsive JWEAU-A10 and the TCD-CD3 inert JWEAU-C6 T cells with compounds known to induce transcriptomic noise by promoting unspecific transcriptional elongation and then stimulated the cells α-CD3/CD28 mAbs. Cells were treated overnight with (**A**) the HDAC inhibitor SAHA (300 nM), (**B**) the cell differentiating agent HMBA (2mM), and (**C**) the BET inhibitor JQ1 (10µM) and then treated with increasing concentrations of α-CD3/CD28 mAbs. HIV-1 reactivation levels were determined by flow cytometry for GFP expression 24 hours post CD3/CD28 activation. Data represent the mean ± standard deviation of at least three independent experiments. The significance of differences between experimental conditions was determined by one-way ANOVA with multiple comparisons.

### Probing the role of individual network nodes in CD3 inertness and HIV-1 latency control

The second possibility would be that individual, highly linked network nodes act as molecular switches that control latent HIV-1 infection, or are key to CD3-inertness, and if targeted alter the stability of latent HIV-1 infection events (Supplemental Figure 7). We would not expect that interference with these key network nodes would directly trigger HIV-1 reactivation but anticipated that pharmacological targeting of the selected network nodes would either boost or abrogate TCR/CD3 activation-induced HIV-1 reactivation. Ideally, interference would restore the ability of TCR/CD3 activation to trigger HIV-1 reactivation in the otherwise inert JWEAU-C6 T cells.

We chose three targets for which selective, clinically relevant inhibitors were available and that were (i) all highly ranked nodes in in the protein-protein interaction network describing the difference between primary T cells from healthy individuals and HIV/ART patients (Table 1) and (ii) central to the network describing the kinomic differences between CD3-responsive and CD3-inert latently HIV-1 infected T cell clones (Table 3). By these means, we would prioritize targets that occur in a cell type independent manner and are less likely to be relevant only in immortalized latently HIV-1 infected T cell clones. Based on these criteria, we chose to target (i) Src (dasatinib), (ii) Raf (sorafenib) and (iii) STAT3 (S31-201). Dasatinib is approved for the treatment of chronic myelogenous leukemia and acute lymphoblastic leukemia. Sorafenib is approved for the treatment of certain kidney and liver cancers and certain forms of Acute Myeloid Leukemia. The selective STAT3 inhibitor S31-201 had been successfully used in several mouse cancer models [110, 111]. We pretreated JWEAU-A10 T cells or long-term HIV-1 infected T cell populations that we enriched for latently HIV-1 infected T cells (see methods section) with increasing concentrations of the three drugs, and, after 24 hours, stimulated with a suboptimal concentration of anti-CD3/CD28 mAbs. Initial use of suboptimal CD3/CD28 stimulation would facilitate the discovery of inhibitory or additive/synergistic effects of the drug on TCR/CD3 signaling. Consistent with the reported central role of Src proteins in TCR/CD3 signaling, inhibition of Src by sub-cytotoxic concentrations of dasatinib effectively abrogated TCR/CD3 triggered HIV-1 reactivation, in both JWEAU-A10 T cells (Figure 6A) and an enriched latently HIV-1 infected T cell population (Figure 6D) [112-115]. It is noted that dasatinib targets several Src family members, which certainly could explain the observed potent effect. While this finding is not surprising, and likely therapeutically irrelevant, it shows the ability of network software to correctly identify and prioritize functionally important protein targets. In contrast to the inhibitory effect of dasatinib, the Raf inhibitor sorafenib boosted the ability of suboptimal CD3/CD28 mAb combinations to trigger HIV-1 reactivation. Sorafenib concentrations as low as 80nM increased the ability of a suboptimal CD3/CD28 stimulation signal to trigger HIV-1 reactivation in JWEAU-A10 T cells from 10% to >20%. A boosting effect could already be observed at lower sorafenib concentrations (Figure 6B). A less potent but significant reactivation-boosting effect was also observed in the latently HIV-1 infected T cell population (Figure 6E). The limited effect of sorafenib on the cell population is another indicator of the heterogeneity between latently HIV-1 infected T cells. Finally, the STAT3 inhibitor S31-201 showed robust enhancement of CD3/CD28 triggered HIV-1 reactivation in JWEAU-A10 T cells (Figure 6C), and importantly, had a strong reactivation-boosting effect in the latently HIV-1 infected T cell populations (Figure 6F).

**Figure 6:**
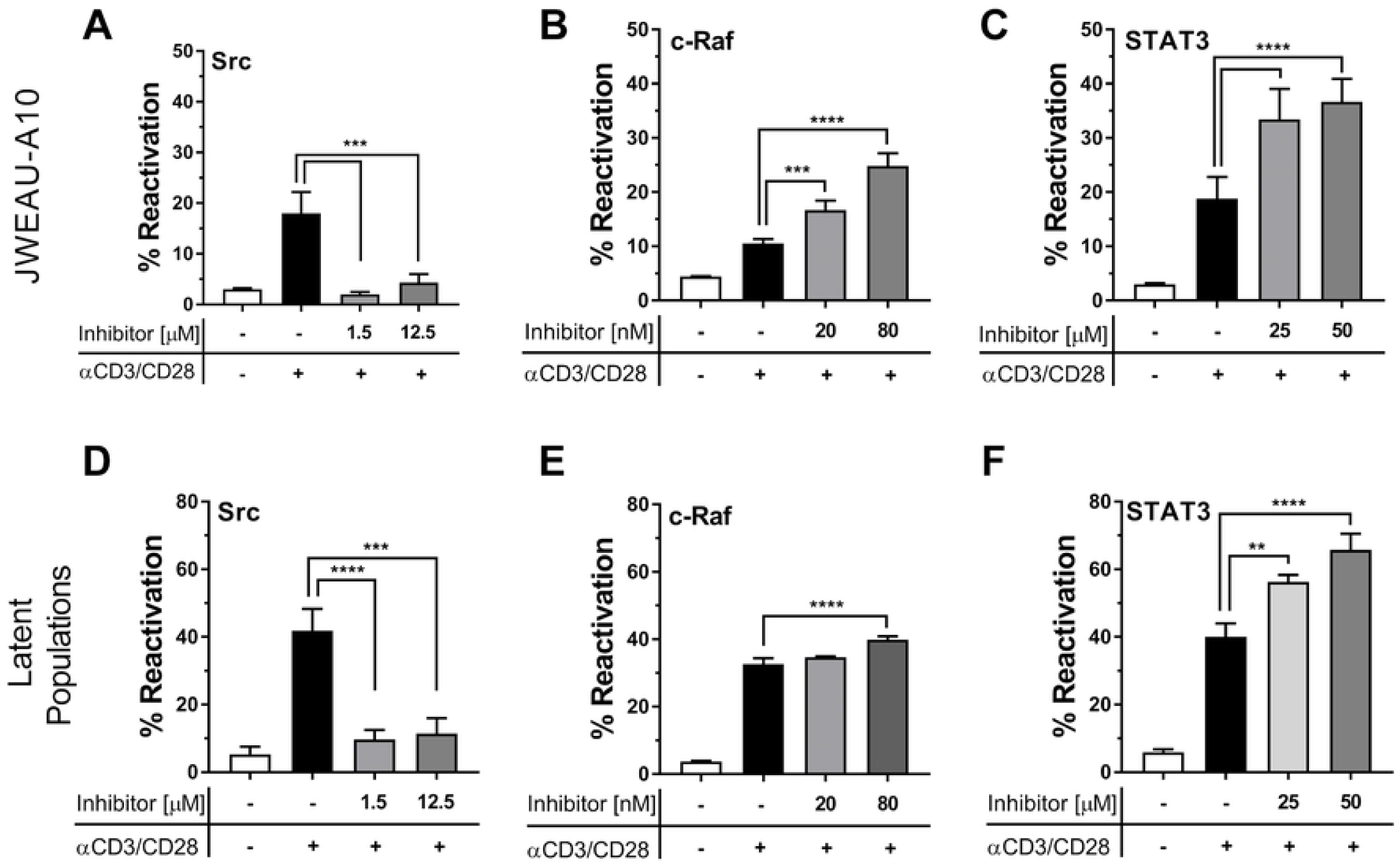
Pharmacological targeting of network hubs alters TCR/CD3 responsiveness of latently HIV-1 infected T cells. To validate that network analysis correctly predicted and prioritized drug targets that affect the ability of TCR/CD3 stimulation to trigger HIV-1 reactivation we tested the effect of inhibitors against Src, Raf and STAT3 on the ability of a-CD3/CD28 mAbs to trigger HIV-1 reactivation. JWEAU-A10 T cells (A-C) or populations enriched for latently HIV-1 infected T cells (D-F) were incubated overnight with different concentrations of the Src inhibitor dasatinib (A, D), the c-Raf inhibitor sorafenib (B, E), or the STAT3 inhibitor S31-201 (C, F) before stimulation with a sub-optimal concentration of α-CD3/CD28 mAbs. Data represent the mean ± standard deviation of at least three independent experiments. The significance of differences between experimental conditions was determined by one-way ANOVA with multiple comparisons.

Having demonstrated that c-Raf and STAT3 inhibition boosted the effect of suboptimal CD3/CD28 stimulation, we next explored whether either c-raf inhibition or STAT3 inhibition would actually reprogram latently HIV-1 infected T cells that are TCR/CD3-inert to become CD3-responsive. This would mean that following either sorafenib or S31-201 treatment, optimal CD3/CD28 stimulation would trigger HIV-1 reactivation in the sub-population of latently HIV-1 infected T cells that are otherwise only responsive to PMA treatment (CD3 inert). As seen in Figure 7A, sorafenib treatment resulted in a small, statistically not significant increase in the percentage of HIV-1 expressing T cells following optimal CD3/CD28 activation, but the majority of the CD3-inert T cell fraction that was revealed by PMA stimulation could still not be addressed. The STAT3 inhibitor S31-201 exhibited a much stronger boosting effect. Pretreatment with S31-201 followed by CD3/CD28 stimulation produced HIV-1 reactivation levels that were similar to the reactivation levels produced by PMA (Figure 7B), indicating that STAT3 inhibition would restore the ability of a large fraction of the initially CD3-inert T cells to generate an effective TCR/CD3 pathway signal. Of note, neither sorafenib nor S31-201 had any effect on the ability of CD3/CD28 mAb combinations to trigger HIV-1 reactivation in the inert JWEAU-C6 T cells, suggesting that extreme CD3-inertness likely cannot be reversed (Supplemental Figure 8).

**Figure 7:**
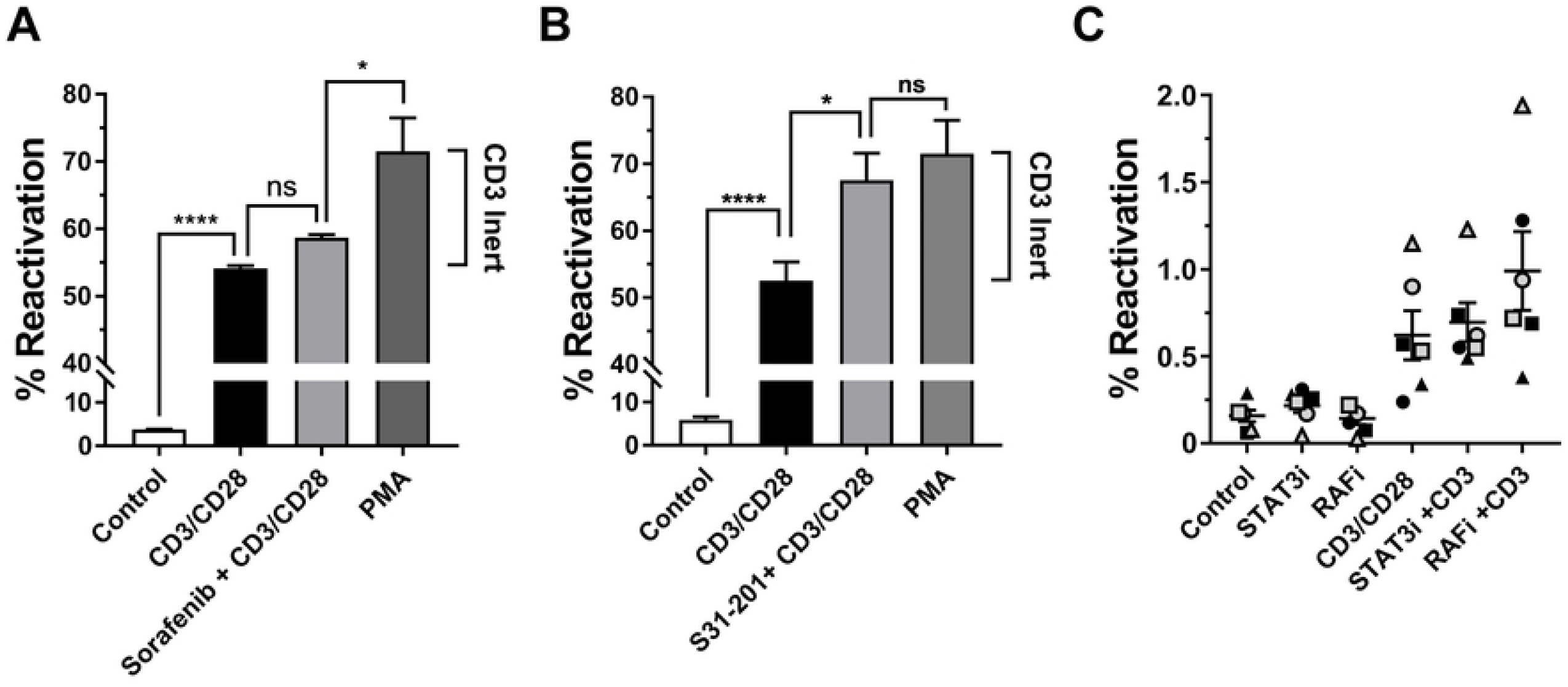
Restoring CD3 responsiveness in latently HIV-1 infected T cells. To test whether pharmacological interventions that were prioritized by network analysis would specifically restore the ability of the CD3 signaling pathway to trigger latent HIV-1 infection in CD3-inert T cells, we stimulated latently HIV-1 infected T cell populations with α-CD3/CD28 mAbs without any pretreatment or following overnight pretreatment with optimal concentrations of (**A**) the c-Raf inhibitor sorafenib or (**B**) the STAT3 inhibitor S31-201 (50 µM). PMA stimulation was used as a positive control to determine the maximum achievable reactivation level in the T cell population. The difference between the reactivation levels accomplishable by α-CD3/CD28 mAb activation and PMA stimulation constitutes the TCR/CD3-inert T cell reservoir (CD3 inert). Any increase of HIV-1 reactivation levels above the ability of α-CD3/CD28 mAbs to trigger HIV-1 reactivation means that the therapeutic intervention allows antibody stimulation to target previously activation inert latently HIV-1 infected T cells. (**C**) Using a HIV-GFP reporter virus latently HIV-1 infected T cell populations were established in primary CD4+ T cell populations obtained from five different donors. The T cells were left unstimulated (control) or treated overnight with optimal concentrations of the STAT3 inhibitor S31-201 or the Raf inhibitor sorafenib. Each of these cultures was then split and left either untreated or stimulated with α-CD3/CD28 mAbs. HIV-1 reactivation was determined by flow cytometry 72 hours post α-CD3/CD28 mAbs addition using GFP expression as a surrogate marker of active HIV-1 expression. The significance of differences between experimental conditions was determined by one-way ANOVA with multiple comparisons.

Finally we tested whether these findings were transferable to latently HIV-1 infected primary T cell models, a possibility that was suggested by the similarity of signaling network changes identified in primary T cells from HIV/ART patients (Table 1) and latently HIV-1 infected T cell lines (Table 3). As for the experiments shown in Figure 2A, we generated latently HIV-1 infected primary T cell cultures using CD4+ T cells from 5 different individuals and tested whether c-raf or STAT3 inhibition here would also boost the ability of CD3/CD28 stimulation to trigger HIV-1 reactivation (Figure 7C). Similar to what was observed in latently HIV-1 infected T cell lines, neither sorafenib nor S31-201 by themselves had any HIV-1 reactivating effect, but both drugs exhibited the ability to boost CD3/CD28 mAb-induced HIV-1 reactivation in a donor-dependent manner. For two of the donors, treatment with both drugs increased CD3/CD28-induced reactivation, whereas for the remaining three donors only one of the drugs provided boosting activity. The observed donor variation is common to models of latent HIV-1 infection that utilize replication competent non-attenuated virus, but it also points out that heterogeneity caused by the immunological history of primary T cells likely adds to the complexity of HIV-1 latency control in primary T cells. The ability of systems biology data that were generated in T cell line models to effectively predict drug targets in latently HIV-1 infected primary T cells provides additional evidence that TCR/CD3 inertness is a fundamental mechanism that may consistently affect T cells that survive the infection.

## DISCUSSION

At the time it is assumed that latent HIV-1 infection events are established as activated T cells become HIV-1 infected before transitioning to a long-lived resting memory phenotype [1, 116-120]. The resting status of memory T cells is thought to deplete HIV-1 of essential transcription factors and thereby restrict its ability to express its genes, forcing the virus into a latent state [121-123]. Based on this concept, it is assumed that T cell activation through antibody-mediated stimulation of the TCR/CD3 complex, the experimental equivalent of cognate antigen recognition, would trigger HIV-1 reactivation. It has by now been conclusively demonstrated that this is correct only for a part of the latent HIV-1 reservoir, whereas a large portion of the latent reservoir remains recalcitrant to TCR/CD3 activation [15, 36]. We thus hypothesized that a part of the HIV-1 reservoir is established in T cells that are either TCR/CD3 activation inert prior to infection or are rendered TCR/CD3 activation resistant by the cellular response to the actual infection event.

In this study we provide formal experimental evidence that TCR/CD3 inertness (i) is generated during HIV-1 infection (Figures 1, 2), (ii) is a fundamental mechanism that is conserved between primary T cells and T cell lines (Figure 2) and (iii) affects a substantial portion of the T cells serving as host cells for the latent HIV-1 reservoir (Figures 2, 7). The existence of such an activation-inert reservoir T cell population would conclusively explain why cognate antigen recognition does not deplete the latent HIV-1 reservoir over time. We further demonstrate that host cells of latent HIV-1 infection events are characterized by extensive stable changes to their signaling networks (Tables 1 and 3). TCR/CD3 inertness was caused by (iv) the occurrence of high-level transcriptomic noise or could be induced by the generation of transcriptomic noise (Figure 5), and (v) was associated with selective changes to the TCR/CD3 signaling pathway, which could be targeted with pharmacological inhibitors to largely reverse TCR/CD3 inertness (Figures 6,7).

The observed presence of transcriptomic noise explains the findings by others that described a high degree of heterogeneity between individual latently HIV-1 infected T cells [16, 17]. Our research extends on these studies in primary T cells and demonstrates that this diversity is not necessarily the result of a pre-existing heterogeneity caused by the different immunological history or differentiation status of T cells that later on become latently HIV-1 infected. While such T cell diversity may add to the observed heterogeneity in primary T cells (Figure 7C), we show that heterogeneity is generated by a fundamental mechanism that must be part of the cellular response to the actual infection event. Heterogeneity between latently HIV-1 infected T cells was observed at the protein and gene expression level (Figure 4, and Tables 1-4). Given that gene expression analysis is now the basis of most studies, it was somewhat surprising that RNA-seq analysis would indicate extensive changes to the gene expression landscape, but network analysis software could not efficiently integrate these data into interaction networks or for that matter assign functional motifs to the data (Supplemental Table 2). Even hierarchical clustering followed by subtree cluster analysis would not directly identify potential molecular switches that were to affect TCR/CD3 mediated HIV-1 reactivation. This was even more surprising as the phenotype, TCR/CD3 inertness, was already predetermined by functional assays (Figures 1-3). In contrast, kinome analysis that directly determined systemwide changes to protein expression or phosphorylation patterns, immediately identified the underlying molecular phenotype (Table 4). The data emphasize the need to investigate HIV-1 latency control mechanisms at the level of both, RNA regulation and protein functionality.

The formal demonstration that heterogeneity at the gene expression and protein expression/phosphorylation level extends to functional diversity has implications for therapeutic strategies that seek to trigger HIV-1 reactivation in patients. The data imply that single drug interventions will be insufficient to accomplish complete viral reactivation and subsequent elimination. TCR/CD3 inertness was not the only functional consequence of the molecular changes that can occur in host cells of latent HIV-1 infected T cells. While we do not detail this phenomenon, we found that TLR pathway signaling can also be impaired (Figure 3), which would have implications for efforts to identify HIV-1 reactivating agents that address the TLR/Myd88 pathway [52-56]. Given that our data detail the dysfunctional nature of latently HIV-1 infected T cells, and the seemingly random nature of observed changes, it needs to be assumed that the further upstream a NF-κB activating pathway is targeted, the less likely it is that an efficient, HIV-1 reactivating NF-κB signal is being triggered. The results thus support the idea that optimal HIV-1 reactivation agents should induce NF-κB activation by targeting proteins as immediate upstream of NF-κB as possible (e.g. directly interfere with the NF-κB/IκB complex), which reduces the likelihood that changes to the host cells affect drug efficacy. Pathways that activate NF-κB, but are essential to the long-term survival of the infected T cell could be most attractive. Here functionality needs to be assumed, as the cells would otherwise be eliminated shortly after the infection event reduces functionality of such an essential pathway. This may make the CD3 or TLR pathways suboptimal targets for LRAs, unless possibly used in a combination therapy. By the same token, the extensive heterogeneity would favor the development of drugs like SMAC mimetics such as AZD5582 that also activate the non-canonical NF-κB pathway. SMAC mimetics, including AZD5582 have already been demonstrated to trigger HIV or SIV reactivation [124-126]. A possible advantage of SMAC mimetics as LRAs could be that they also target an important survival pathway for cells. SMAC mimetics imitate the activity of a protein call Second Mitochondrial-derived Activator of Caspases, a pro-apoptotic mitochondrial protein that is an endogenous inhibitor of a family of cellular proteins called the Inhibitor of Apoptosis Proteins (IAPs), which in turn are important regulators of cell death and survival. The importance of such a pathway for long-term survival of the cells should make it less likely that this pathway is impaired in host cells of latent HIV-1 infection events. Functional conservation of this pathway would increase the likelihood of SMAC mimetics to broadly trigger HIV-1 reactivation within the population of latently HIV-1 infected T cells, despite the heterogeneity and pre-existing inertness in other pathways.

Conceptually, the provision of experimental evidence that many host cells of latent HIV-1 infection events are extensively altered in their signaling networks and gene expression patterns and in extension, are altered in many of their functionalities, raises the possibility to affect the stability of latent HIV-1 infection events in these cells through an alternative approach. Therapeutic interventions that would restore host cell functionality may enable TCR/CD3 activation-inert latently HIV-1 infected T cells to properly respond to cognate antigen recognition, and the ensuing cellular increase in NF-κB activation would be sufficient to trigger HIV1 reactivation. Such a scenario would focus on the development of cellular reprograming strategies and remove the requirement for a potent, systemwide, stimulatory therapeutic intervention, which would here be provided by a natural process, specific antigen recognition. Given the general impairment of T cell functionality in PBMCs from HIV/ART patients that we observed (Figure 1), a T cell status that seems to have many signaling network similarities with latently HIV-1 infected T cells, such a strategy may also improve the overall immune status of HIV-1 patients.

## MATERIAL AND METHODS

### Cell culture and reagents

All T cell lines were maintained in RPMI 1640 supplemented with 2 mM L-glutamine, 100 U/ml penicillin, 100 µg/ml streptomycin and 10% heat inactivated fetal bovine serum. Fetal bovine serum (FBS) was obtained from HyClone (Logan, Utah) and was tested on a panel of latently infected cells to assure that the utilized FBS batch did not spontaneously trigger HIV-1 reactivation [48, 127]. The phorbol esters, Phorbol 12-myristate 13-acetate (PMA), prostratin, along with HMBA, Flagellin, and Trichostatin A (TSA) were purchased from Sigma. Recombinant human TNF-α a well established trigger of HIV-1 reactivation was purchased from Gibco. The PKC activator Bryostatin and Sodium Butyrate (NaBu) were purchased from EMD Millipore. Vorinostat (SAHA) was purchased from Selleck chemicals. Specific Inhibitors or recombinant proteins such as the STAT3 inhibitor S31-201 (NSC 74859), the Src inhibitor dasatinib, or the c-Raf inhibitor sorafenib were all purchased from Fisher Scientific. Anti-CD3/CD28 antibody-coated beads (ImmunoCult) were purchased from STEMCELL technologies (Vancouver, CA).

### Human Sample disclosure and Information

Healthy, HIV-seronegative adults and HIV-seropositive subjects were recruited from the 1917 Clinic cohort at the University of Alabama at Birmingham to donate peripheral blood. All HIV-1 seropositive subjects were on ART and with undetectable viral loads (VL <50 copies/mL) for a median of 10 months (6.5-17.3 months). In accordance with the specific protocol for this project that was approved by the UAB Committee on Human Research (IRB-160715008), all subjects provided written informed consent for all biologic specimens and clinical data used in this study.

### Model of HIV-1 latency in primary T cells

CD4+ T-cells were isolated from PBMCs from healthy donors using a negative CD4+ T cell isolation kit (Stemcell), and were then rested overnight in RPMI (10% FBS, 1% Pen/Strep, 1% L-Glutamine, and IL-2 (10 IU/ml). The T cells were stimulated with Immunocult (Stemcell) at a concentration of ∼2 beads/cell and incubated at 37° for 3 days. On day three post stimulation the T cells were placed in fresh RPMI medium at a density of 5 million cells/well in a 6 well plate and infected with a HIV-GFP reporter virus (pBR43IeG-NA7nef) [74]. After infection cells were pelleted by centrifugation and placed in fresh RPMI medium. After 48 hours GFP levels were determined as surrogate marker of HIV-1 infection using flow cytometry to confirm successful infection. To prevent further viral replication 3TC and EFV and were added and the cells were cultured for 30 days. Medium was replenished twice a week. After 30 days the T cell cultures were sorted using a BD Aria cell sorter to remove all remaining GFP+, and therefore actively infected T cells to reduce the signal background. Cells were then plated at a density of 5×10^5^ to 1×10^6^/ml into the described experimental conditions. 3 days post reactivation the cell cultures were analyzed for the frequency of GFP+ T cells using a BD LSRII flow cytometer acquiring at least 100,000 events per experimental condition.

### Generation of latently HIV-1 infected T cell populations and T cell clones

The latently HIV-1 infected T cell populations were generated by infecting a Jurkat T cell-based reporter cell population (JR5D4 cells) [50] with the primary HIV-1 patient isolate WEAU [51] or the classic laboratory strain HIV-1 NL43. Starting two days post infection a reverse transcriptase inhibitor (3T3) was continuously added to prevent viral replication and the establishment of pre-integration latency. On day 4 post infection we selected cultures with infection levels around 40% and maintained these until day ∼60 when active infection levels had mostly declined to levels between 1-5%. In the HIV-1 WEAU infected T cell population, addition of PMA to this T cell population revealed the presence of a reactivatable HIV-1 reservoir that varied in size between 4-10%. The HIV-1 WEAU infected bulk cell populations were plated in a limiting dilution and tested for CD3 inertness and PMA responsiveness, to generate the differentially responsive cell clones JWEAU-A10 (responsive) and JWEAU-C6 (inert). To generate T cell populations that were enriched for latently HIV-1 infected T cells, we exploited the reported ability of host cells of latent HIV-1 infection events to repeatedly shut down the virus into a latent state. HIV-1 reactivation was triggered using PMA and on day 2 post activation cell sorting was performed for GFP+ T cells. Over a period of 3-4 weeks, active HIV-1 infection in these cells was again shut down into a latent state, resulting in an enriched population in which ∼80% of the cells would hold reactivatable, latent HIV-1 infection events.

### CFSE Proliferation assays

PBMCs were resuspended in PBS at roughly 10^7^ cells/ml and labeled at room temperature with 5,6-carboxyflourescein diacetate succinimidyl ester (CFSE; Molecular Probes, Eugene, OR) at 1.25μM for 4 mins. The cells were washed twice in PBS (10% FBS). Cells were then resuspended in complete media (RPMI + 10% AB sera, 50 U/ml of penicillin/streptomycin, 2mM L-glutamine). Cells were stimulated with α-CD3/CD28 mAbs for 4 days, after which proliferation was measured by loss of CFSE fluorescence using a BD LSRII flow cytometer with >100,000 lymphocyte events collected per sample. Unstimulated cells were used for negative controls.

### Kinexus™ Antibody Microarray-based analysis

Kinome analysis to study protein expression and phosphorylation levels was done using Kinex™ microarray analysis. 50 μg of cell lysate protein (∼5×10^6^ cells) from each sample or experimental condition were covalently labeled with a proprietary fluorescent dye according to the manufacturer’s instructions (Kinexus, Canada). After the completion of the labeling reaction, any free dye was removed by gel filtration. After blocking non-specific binding sites on the array, an incubation chamber was mounted onto the microarray to permit the loading of two side by side samples on the same chip. Following sample incubation, unbound proteins were washed away. KAM-850 arrays detect 189 protein kinases, 31 protein phosphatases and 142 regulatory subunits of these enzymes and other cell signaling proteins. This array provided information on the phosphorylation state of 128 unique sites in protein kinases, 4 sites in protein phosphatases and 155 sites in other cell signaling proteins. KAM-900 chips are spotted in duplicates with over 870 antibodies. 265 pan-specific antibodies and 613 phosphosite specific antibodies. Each array produced a pair of 16-bit images, which are captured with a Perkin-Elmer ScanArray Reader laser array scanner (Waltham, MA). Signal quantification was performed with ImaGene 8.0 from BioDiscovery (El Segundo, CA) with predetermined settings for spot segmentation and background correction. The background-corrected raw intensity data were logarithmically transformed with base 2. Since Z normalization in general displays greater stability as a result of examining where each signal falls in the overall distribution of values within a given sample, as opposed to adjusting all of the signals in a sample by a single common value, Z scores are calculated by subtracting the overall average intensity of all spots within a sample from the raw intensity for each spot, and dividing it by the standard deviations (SD) of all of the measured intensities within each sample [128]. Z’ ratios were further calculated by taking the difference between the averages of the observed protein Z scores and dividing by the SD of all of the differences for that particular comparison. Calculated Z’ ratios have the advantage that they can be used in multiple comparisons without further reference to the individual conditional standard deviations by which they were derived.

### RNA-seq analysis

Total RNA was extracted using the Qiagen RNeasy mini kit. Genewiz (Plainfield, NJ) prepared cDNA libraries and performed sequencing. For RNA-seq data processing and analysis, raw paired reads were first adapter-trimmed from fastq files using TrimGalore! (http://www.bioinformatics.babraham.ac.uk/projects/trim_galore/). Reads were aligned to Hg19 using STAR [129], and count matrices were generated using HTSeq-count [130]. DESeq2 was used to normalize read counts (CPM) and analyze differential expression [131]. For the analysis of global transcriptional regulation, genes were considered differentially expressed if they had an adjusted *P*-value <0.01 by the likelihood ratio test. For pairwise comparisons, genes were considered differentially expressed if they had at least a 1.5-fold change, adjusted *P*-value < 0.01, and at least one signal > 250 CPM, as qPCR validation repeatedly failed to confirm lower signals. Pheatmap (*) was used to row-normalize, cluster (via hclust, using the complete linkage method), and visualize global transcriptional changes. Gene Ontology analysis was conducted using Metascape [132].

### Network Analysis

MetaCore software (Thomson Reuter) was used to generate shortest pathway interaction networks that are presented. Pathway specific GO filters or tissue specific filters were used to prioritize nodes and edges. MetaCore was further used to identify pathway associations of the identified input dataset, using network enrichment analysis.

### Statistical Analysis

Statistical significance of differences between multiple experimental conditions was determined by ANOVA with multiple comparisons (Tukey’s correction) (*p<0.05, **p<0.01, ***p<0.001, ****p<0.0001).

## FIGURE LEGENDS

**Supplemental Figure 1: CD4+ T cells from HIV/ART patients exhibit an altered proliferation response following CD3/TCR stimulation.** Proliferation of primary CD4+ T cells from **(A)** HIV/ART patients or **(B)** healthy donors measured using CFSE staining reveals a distinct pattern of cellular division representative plots shown (n=11).

**Supplemental Figure 2: Network analysis of kinome data obtained from T cells from HIV/ART patients.** Protein interaction networks were generated using T cells from HIV/ART patients in comparison to T cells from healthy donors. The direct protein-protein interaction network was generated using MetaCore software and describes the signal transduction pathways that are altered in HIV/ART patients.

**Supplemental Figure 3: TNF-α triggered HIV-1 reactivation in the latently HIV-1 infected T cell clones.** JWEAU-A10 or JWEAU-C6 T cells were stimulated with increasing amounts of TNF-a. HIV-1 reactivation levels were determined by flow cytometric analysis, using GFP expression as a surrogate marker of active HIV-1 expression. Data represent the mean ± standard deviation of at least three independent experiments.

**Supplemental Figure 4: Transcriptomic analysis shows global changes to transcriptional regulation in latently infected cells.** Significantly altered RNA signals were used to generate protein-protein interaction networks using the MetaCore direct interaction algorithm. The network visualizes the large amount of unconnected seed nodes, that were likely regulated by bystander effects.

**Supplemental Figure 5: Gene Ontology analysis of distinct clusters within RNA-seq data.** Gene ontology analysis of the individual gene clusters in Figure 4C.

**Supplemental Figure 6: Focused analysis of genes involved in TCR/CD3 mediated T cell activation.** Expression levels of genes involved in T cell activation were compared between Jurkat T cells and the two latently HIV-1 infected T cell lines, JWEAU-A10 and JWEAU-C6. The gene expression heat map visualizes a general down regulation of genes involved in T cell activation in latently HIV-1 infected T cell clones, but no relevant differences between the TCR/CD3 activation responsive JWEAU-A10 T cells and the activation inert JWEAU-C6 T cells.

**Supplemental Figure 7: Kinome analysis of differentially responsive cell lines generates highly connected protein interaction networks.** Protein Interaction networks were generated from significantly altered protein signals between JWEAU-A10 and JWEAU-C6s on the Kinexus kinome chip as previously described.

**Supplemental Figure 8: Protein targets fail to rescue completely CD3 inert latently HIV-1 infected cells:** To determine if the inhibitors against Src, Raf and STAT3 were capable of rescuing completely CD3 inert latently infected T-cells compounds were titrated on JWEAU-C6 cells prior to overnight stimulation with αCD3/CD28 mABs. Data represent the mean ± standard deviation of at least three independent experiments.

